# Distributed and dynamic intracellular organization of extracellular information

**DOI:** 10.1101/192039

**Authors:** Alejandro A. Granados, Julian M. J. Pietsch, Sarah A. Cepeda-Humerez, Gašsper Tkačik, Peter S. Swain

## Abstract

Although cells respond specifically to environments, how environmental identity is encoded intracellularly is not understood. Here we study this organization of information in budding yeast by estimating the mutual information between environmental transitions and the dynamics of nuclear translocation for ten transcription factors. Our method of estimation is general, scalable, and based on decoding from single cells. The dynamics of the transcription factors are necessary to encode the highest amounts of extracellular information, and we show that information is transduced through two channels: generalists (Msn2/4, Tod6/Dot6, Maf1, and Sfp1) can encode the nature of multiple stresses but only if stress is high; specialists (Hog1, Yap1, and Mig1/2) encode one particular stress, but do so more quickly and for a wider range of magnitudes. Each transcription factor reports differently, and it is only their collective response that distinguishes between multiple environmental states. Changes in the dynamics of the localization of transcription factors thus constitute a precise, distributed internal representation of extracellular change, and we predict that such multi-dimensional representations are common in cellular decision-making.

**Significance Statement:** To thrive in diverse environments, cells must represent extracellular change intracellularly despite stochastic biochemistry. Here we introduce a quantitative framework for investigating the organization of information within a cell. Combining single-cell measurements of intracellular dynamics with a new, scalable methodology for estimating mutual information between time-series and a discrete input, we demonstrate that extracellular information is encoded in the dynamics of the nuclear localization of transcription factors and that information is lost with alternative static statistics. Any one transcription factor is usually insufficient, but the collective dynamics of multiple transcription factors can represent complex extracellular change. We therefore show that a cell’s internal representation of its environment can be both distributed across diverse proteins and dynamically encoded.

---

All organisms sense their environment and the information gained must be represented internally to elicit a change in behaviour (*1*). Much is understood about such representations in neural systems (*2*). Cells must perform an analogous task (*1*, *3*), encoding intracellularly the information about extracellular environments. Yet little is known about the nature of such encoding.

The activation of transcription factors is thought to provide an internal representation of the environment (*4– 10*), but how information is encoded dynamically, whether information is spread across multiple factors, and how information is read out downstream all remain unclear (Fig. 1A). We do know that the biochemical implementation of such representations is likely to be substantially stochastic (*11*) and that the same biochemistry may be used to sense disparate environments. Furthermore, cells typically have just ‘one shot’ at mounting the appropriate response from these internal representations, with competition being unforgiving for those that delay, at least among microbes (*12–14*). Here we use information theory to investigate how eukaryotic cells answer these challenges through their internal organization of extracellular information.

To do so, we turn to budding yeast and to environmental changes for which we expect information encoding to be key: stresses that compromise growth and evoke adaptive gene expression (*15*). In yeast, extracellular changes are sensed by signaling networks, homologous to networks in higher organisms (*16*), that regulate the activity of transcription factors, often by their translocation either into or out of the nucleus (*16*), analogous to p53 and NF-*κ*B in mammalian cells (*17*, *18*). We therefore consider the movement of these transcription factors as a cell’s internal representation of an environmental transition (Fig. 1B & Fig. S1). The translocations are dynamic and stochastic, and the information available from the full time series of the response could be substantially higher than that available from any temporal snapshot (*9*) (Fig. 1A).

## Results and Discussion

To quantify the information available to the cell, we developed a general and, unlike previous approaches, scalable methodology to estimate the mutual information between the time series of cellular responses and the state of the extracellular environment (SI Appendix; Fig. 1C & Figs. S3–S7). Mutual information, a robust measure of statistical dependency (*19*), allows us both to capture the effects of biochemical stochasticity, which can drive individual responses far from the average, and to avoid *a priori* assumptions about which features, such as the magnitude and frequency, of the response are relevant.

Our method uses techniques from machine learning to train a classifier to predict the state of the environment from the time series of as few as 100 cells (Fig. S3). The classifier output on the testing data can then be used to estimate the mutual information. Formally, this approach provides a lower bound on the true information (SI Appendix), but, biologically, we are quantifying the information that a cell could plausibly recover and act upon after observing a single time series of its response. Although it is unnecessary, we assume that each potential environment is equally probable for simplicity. By varying the duration of the time series used by the classifier, we can determine how quickly cells accumulate information (Fig. 1D). In addition, errors made by the classifier indicate environments that are likely to be confused, giving insight into tasks potentially challenging for the cell.

Tens of transcription factors translocate in yeast (*16*, *20*) and we focus on a representative subset, comprising three classes (*16*) (Fig. 2A): the environmental stress response (Msn2 and its paralog Msn4); specialists, which respond to a particular stress (Mig1 and its paralog Mig2, Hog1, and Yap1); and growth-related transcription factors, which either directly or indirectly regulate the rate of translation (Dot6 and its paralog Tod6, Sfp1, and Maf1). Some of these factors (Msn2/4, Mig1/2, and Dot6/Tod6) have pulsatile dynamics, with stochastic bursts of nuclear localization even in the absence of stress (*20*).

We consider environmental shifts from rich media (2% glucose) into either carbon stress (lower glucose), hyperosmotic, or oxidative stress. Using fluorescent tagging and microfluidics (*21*), we measure the degree of nuclear localization of the transcription factors in hundreds of single cells both before and after the stress is applied (Fig. 2B). The mutual information we calculate then addresses whether an environmental transition can be detected from a typical time series of nuclear localization.

Specialists perform almost optimally in their cognate stress with near the maximum possible information of 1 bit (Fig. 2C). For a single environmental transition, the mutual information is a number between 0 bits (indicating no detectable statistical differences between the dynamics of localization before and after the transition) and 1 bit (the dynamics of localization before the transition is distinct from the dynamics after the transition). We note that different transcription factors encode information in different ways. For example, Dot6 and Sfp1 display different dynamics (Fig. 2B), yet encode the same amount of bits (Fig. 2C). Paralogs, however, do not necessarily carry the same information (compare Tod6 and Dot6).

Information can be encoded within minutes of the environmental transition, and the speed of encoding typically increases the more information is encoded. Defining the encoding delay as the time for the mutual information to reach 50% of its maximum value, information then plateaus earliest in the time series of Mig1/2 (Fig. 2D). Fast responders are therefore typically more accurate, at least for the large stresses considered here.

**Figure 1.**
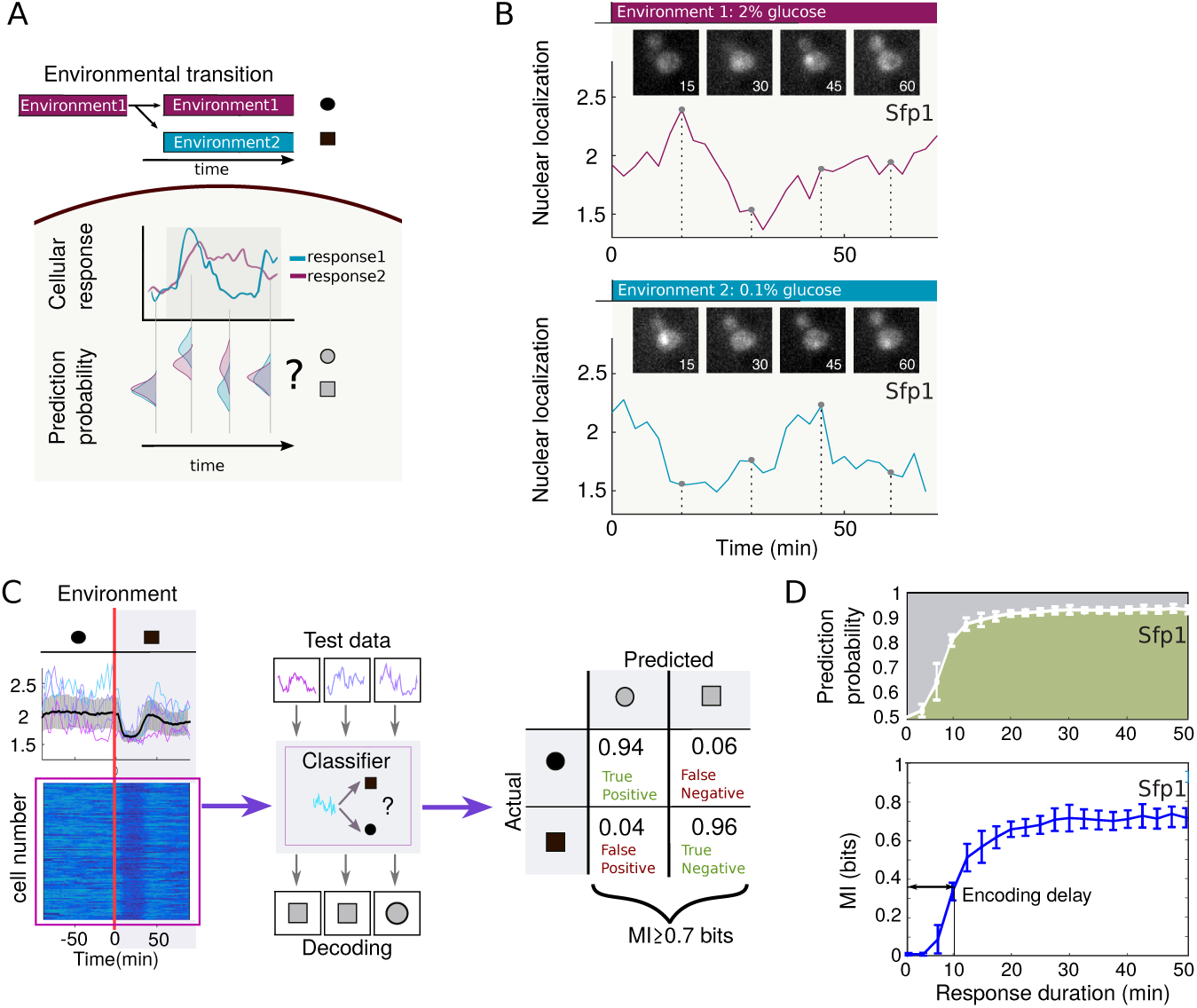
Intracellular responses carry information on extracellular change. (*A*) Environmental transitions trigger dynamic intracellular responses (purple and blue time series), which are often the only indications to the cell that the environment has altered. Biochemical stochasticity can make intracellular responses a poor readout of the new environment, and a probabilistic representation of the possible environmental state based on the instantaneous intracellular response may vary over time. (*B*) Our approach is based on comparing single-cell time series of nuclear translocation (Sfp1, an activator of ribosome biogenesis, is shown) in different environments. The transcription factors are tagged with Green Fluorescent Protein and their dynamics followed in the ALCATRAS microfluidic device (inset & Fig. S2). (*C*) We estimate the mutual information between the time series and the environment by using 70% of the data to train a classifier to classify the time series into the two environments (each coloured line is a single cell with greater nuclear localization indicated by lighter blue). With the remaining data, we determine the confusion matrix (the probability of correctly and incorrectly identifying the environment from a time series) and, from this matrix, a lower bound on the mutual information. (*D*) By increasing the length of the time series used by the classifier (each time series has the same initial point), we estimate both the probability of correct and incorrect predictions and the mutual information as a function of the duration of the response. Here the actual environment is 0.1% glucose (green area indicates the fraction of correctly classified time series). At *t* = 0, the two environments cannot be distinguished, and the best prediction is a random guess, which has a prediction probability of 0.5 and corresponds to 0 bits of information. Error bars are S.D. across biological replicates (*n* = 6).

These general observations hold too for transitions into osmotic and oxidative stress (Fig. 2E & 2F), establishing a hierarchy: in terms of the information and encoding delays, specialists are followed by the environmental stress response, which in turn is followed by the growth-related factors. The details of this hierarchy, however, are stress-specific, indicating that the dynamics of the transcription factors may encode not only the presence but also the nature of the environmental transition.

We therefore extended our method to calculate the mutual information between a single-cell time series and the four environmental states investigated: rich media (before the environmental transition) and the three stresses (after the environmental transition). Not only do we estimate the mutual information (Fig. 3A), but we can also predict how a typical time series is likely to be classified as a function of the duration of the environment (Fig. 3B & Fig. S9)

Although no single transcription factor reaches the maximum of 2 bits, the time series of Msn2, Msn4 and Dot6, carry sufficient information to identify three environmental categories (for example, two environmental states and the remaining two states lumped into a single category) (Fig. 3A & SI Appendix). We observe, however, that the information is instead ‘spread’ so that all environmental states are classified with a greater than 70% accuracy, at least at late times (Fig. 3B: Dot6). In contrast, an ideal specialist should perfectly discriminate one environmental state and lump together the remaining states to encode 0.8 bits (SI Appendix). Indeed, Hog1 and Yap1 do encode this much information (Fig. 3A), and their signalling networks must operate nearly optimally in these high stresses. After only a few minutes, specialists unequivocally identify their associated stress and never report false positives (Fig. 3B: Yap1 & Mig1).

**Figure 2.**
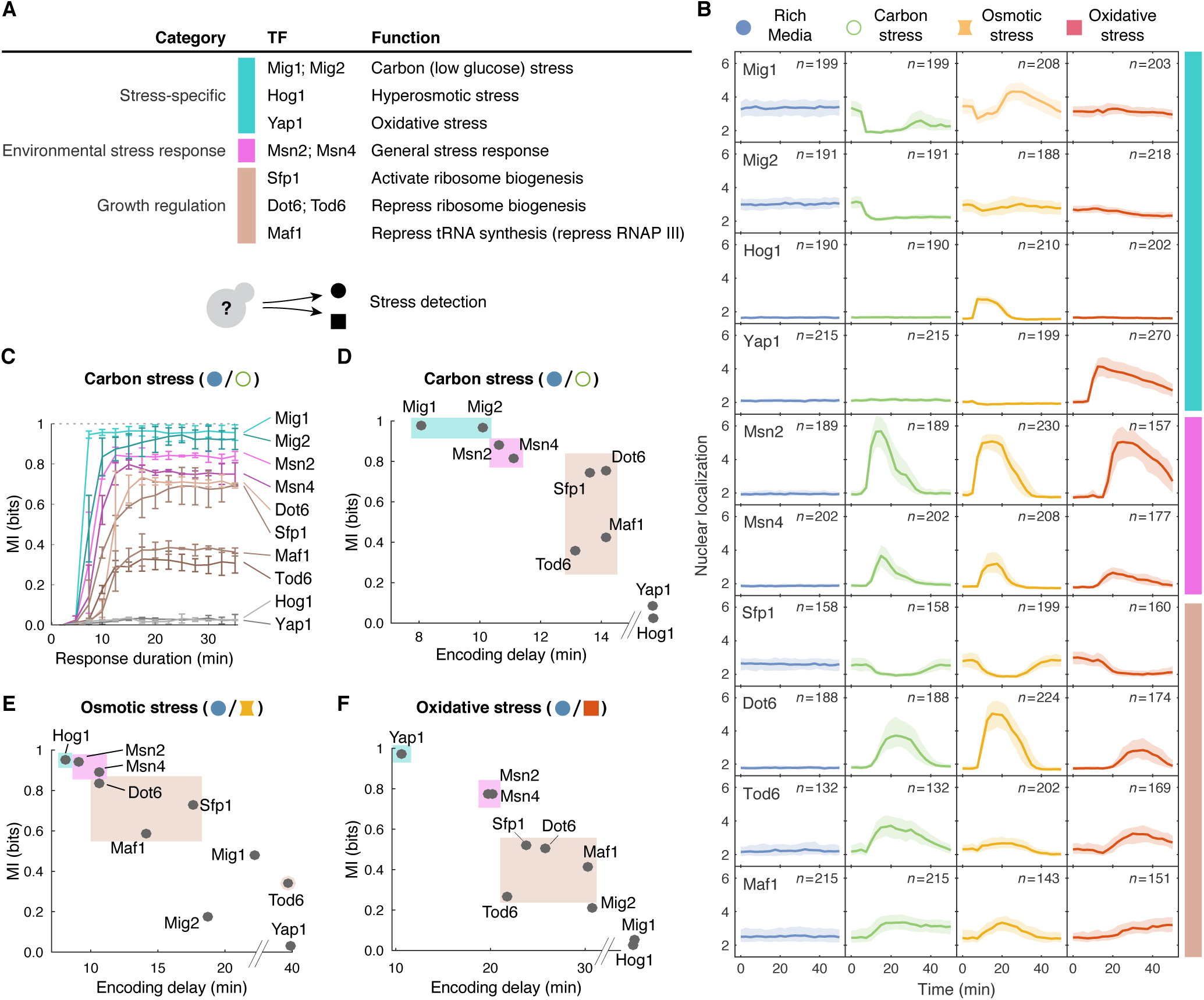
A hierarchy of information encoding, both in bits and encoding delays, holds across environmental transitions. (A) We focus on 10 transcription factors (Hog1 is also a p38-like MAP kinase) and (*B*) quantify nuclear localization across 4 environments: no stress (2% glucose) before the transition (applied as a step-change at *t* = 0) and carbon stress (0.1% glucose), osmotic stress (0.4M NaCl), or oxidative stress (0.5mM H_2_O_2_) afterwards. The median and interquartile range for a representative experiment are shown. (*C*) In response to carbon stress, the mutual information shows a hierarchy for both the number of bits and the speed of the encoding. The maximum possible information is 1 bit (dotted line). (*D, E, F*) The hierarchy’s order is maintained for transitions into other stresses: specialists (blue) encode the most information and are fastest, followed by the environmental stress response (pink), and then the growth-related factors (brown). Mean of 2 experiments per transcription factor.

Conditioning the mutual information on the identity of the environmental states delineates specialists from the other transcription factors (Fig. 3A & Fig. S8), which we term generalists because they encode information on multiple types of stress. Nevertheless, these groups are not mutually exclusive: Mig1 is not only a specialist for carbon stress, but also carries information on the other states at late times, particularly osmotic stress, and has a probability of over 50% of identifying the correct environment, more than twice the 25% probability of a random choice.

**Figure 3.**
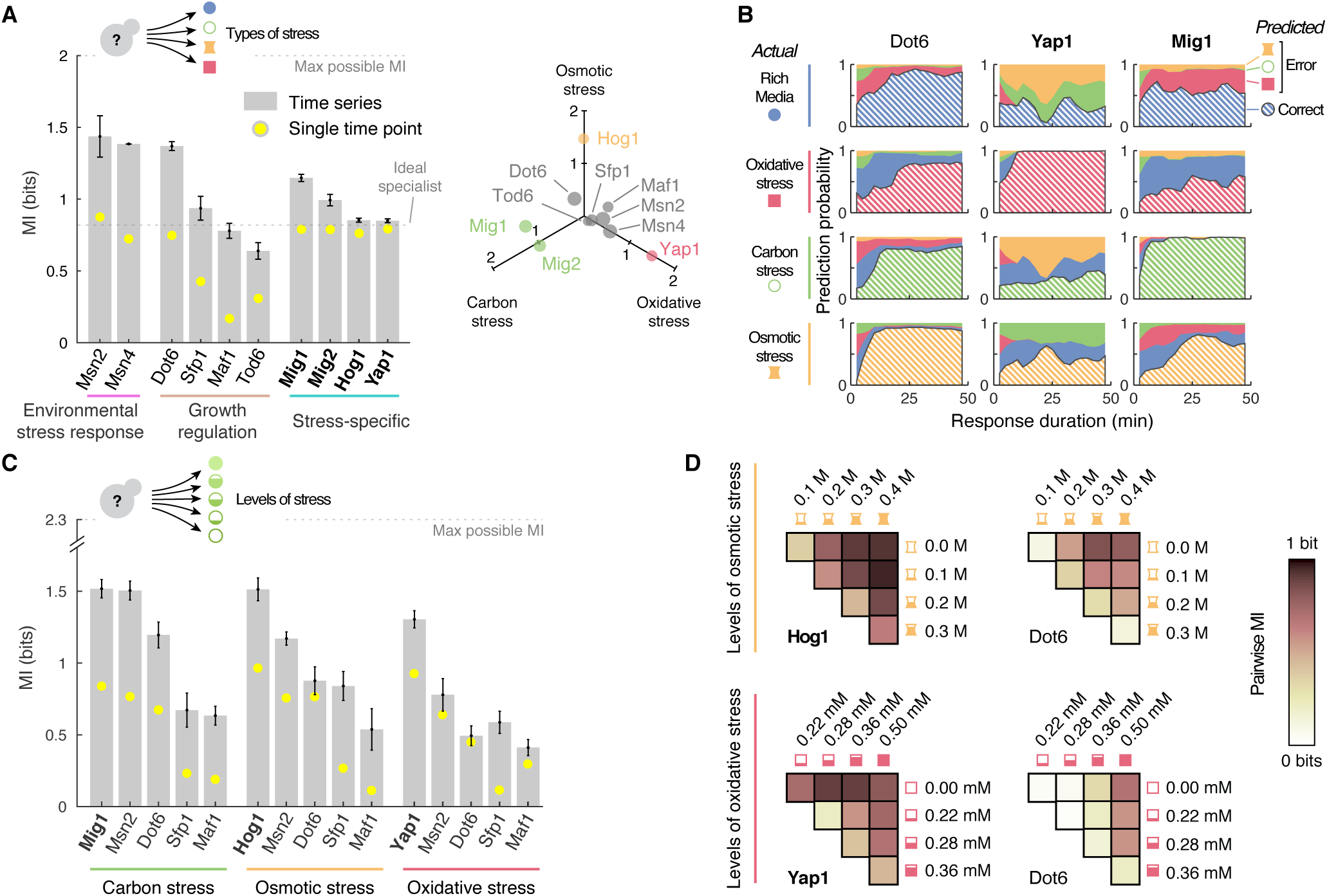
Generalists can identify the nature of large environmental transitions but only specialists distinguish smaller changes. (*A*) The maximum mutual information between a time series of translocation and four environmental states: rich media (before the transition) and carbon stress, osmotic stress, or oxidative stress (after the transition). The mutual information calculated using the best single time point is always lower (yellow dots). Mean and SD for each transcription factor is shown (calculated from 2 independent data sets: 6 experiments per transcription factor). Inset: Conditioning the mutual information on particular environment states differentiates the specialists (in colour) from the others. (*B*) Our methodology estimates the typical probability of inferring each environmental state from a time series. The true state is marked with hash lines. Inference generally improves with the duration of the new environment, and rich media and oxidative stress are confused most often. (*C*) The maximum mutual information between a time series of translocation and 4 levels of stress (after the transition) and rich media (before the transition). Specialists perform best and encode with their entire time series (compare yellow dots with those in **A**). (*D*) Comparing the mutual information between all pairs of environments with the same type of stress at different levels, generalists can identify high stress, but poorly distinguish adjacent levels compared to specialists.

In such high stresses, specialists appear unnecessary because the generalists identify stress so well, but this situation changes if we consider transitions into stresses of lower magnitude (Fig. 3C & Fig. S10). From rich media, we applied four different levels of stress and estimated the mutual information between the time series of translocation and five environmental states: the four levels of stress after the transition and rich media before the transition. Considering the mutual information between the time series and all pairs of the different levels of stress (Fig. 3D & Fig. S12), we see that distinguishing between adjacent levels is most challenging, even for specialists (generalists, such as Dot6, can confuse the low levels of stress and only identify high stress).

Generalists and specialists also differently encode information: generalists usually encode information in their dynamics whereas specialists only do so to distinguish stresses of lower magnitude. By calculating the mutual information between summary features of the single-cell time series and the environmental state (Figs. S12 and S13), we found that the amplitude of a specialist’s initial translocation can suffice to identify its associated stress if that stress is sufficiently severe (yellow dots in Fig. 3A), explaining specialists’ short encoding delays. For transitions into stresses of lower magnitude, however, information is encoded in the dynamics of the specialists’ response (yellow dots in Fig. 3C). Generalists can encode twice the amount of information in their dynamics compared to the best single time point, and both the timing of their initial translocation, particularly for Msn2 and Dot6, and the shape of the times series can be important (SI Appendix).

**Figure 4.**
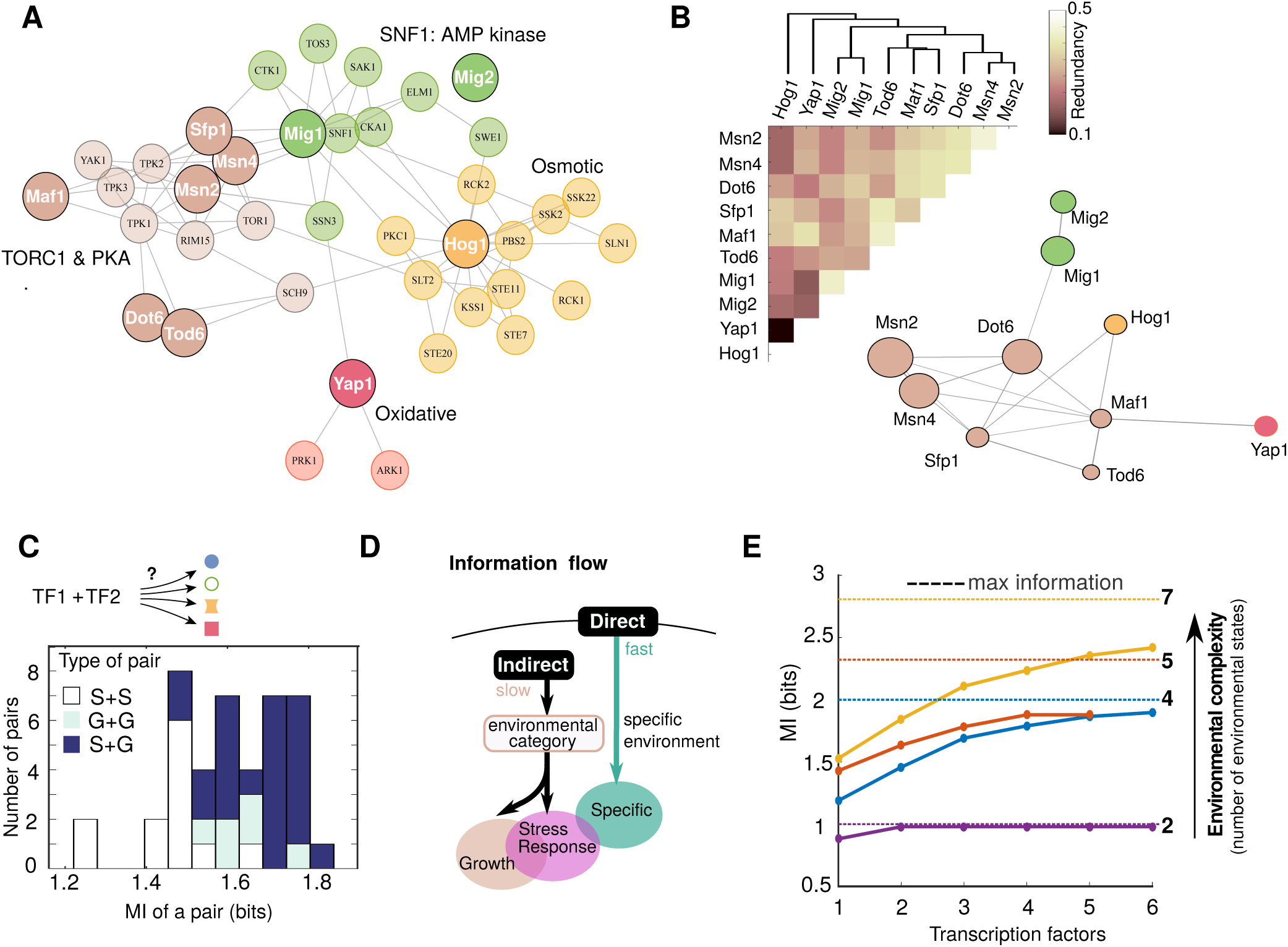
Collectively, transcription factors provide an internal representation of complex environments. (*A*) In the intracellular signaling network (SI Appendix), generalists are co-located and specialists are distinct. Edges are known interactions between either two kinases or a kinase and a transcription factor. Clusters of highly interconnected components have different colors and we include only the transcription factors in our study. (*B*) The redundancy in information between pairs of transcription factors reflects the structure of the signaling network. Edges are proportional to a pair’s redundancy: 1 *-* MI12*/*(MI1 + MI2) (*22*). Inset: the matrix of redundancy for all pairs calculated from an average of 2 datasets (6 experiments per transcription factor). (*C*) A generalist and a specialist typically encode the most information out of all possible pairs of transcription factors. For each pair, time series from one transcription factor were concatenated randomly with the time series of another to calculate the mutual information. Legend: S – specialist; G – generalist. (*D*) Cells have two types of channel to transduce information. The direct channels (specialists) respond to particular extracellular change; the indirect channels (generalists) respond to intracellular changes and so to different categories of extracellular change. (*E*) Only through combinations of transcription factors can information on complex environments be encoded. Different colors represent different numbers of environmental states: orange – 7 states with different types and levels of stress (SI Appendix); red – 5 states with one type of stress at different levels; blue – 4 states (rich media before the transition and 3 types of stress of high magnitude after the transition); purple – 2 states either rich media (before the transition) or stress (after the transition). Plotted lines are the 80’th percentile calculated from all combinations of concatenated time series for a given number of transcription factors (SI Appendix).

To better understand the behavior of generalists and specialists, we asked how the transcription factors are organized within the cell’s networks of signal transduction. Using data on the substrates of kinases (*23*), we confirm (*16*) that Msn2/4 and the growth-related transcription factors are either directly or indirectly targets of protein kinase A (which has isoforms Tpk1-3) and TORC1 or its downstream kinase Sch9 (mammalian S6 kinase) (Fig. 4A). The generalists therefore respond to the cell’s potential for growth: protein kinase A orchestrates the cell’s response to the availability of glucose (*16*) and TORC1 controls the response to the availability of nitrogen (*16*). Similarly, Mig1 is sensitive to the levels of cellular energy through its phosphorylation by AMP kinase (*16*). In contrast, the specialists Hog1 and Yap1 are mostly embedded in their own signalling networks.

The signaling network’s modular architecture is reflected in the organization of information. To understand how information is distributed, we quantified the redundancy between pairs of transcription factors: if regulated by the same upstream signalling, a pair of transcription factors may be completely redundant so that when paired together the amount of information available on the environment does not increase from that available from one of the factors alone. By concatenating two time series, we can estimate the mutual information from simultaneously observing two transcription factors and so quantify their degree of redundancy. Plotting the redundancy (Fig. 4B & Fig. S14), we see a network similar to the network of signal transduction: Msn2/4 and the growth-related factors are together in a core from which the specialists are distinct. The redundancies imply that pairing a generalist with a specialist is best and, indeed, such pairs typically encode the highest information (Fig. 4C). With its distinct signal transduction (Fig. 4A), a specialist can identify the environmental state that is most poorly distinguished by a generalist. For example, Msn2 is best paired with Hog1, and Dot6 is best paired with Yap1 (Fig. S14A).

Despite the three classes of transcription factors, information is transduced through only two channels: specialists and generalists (Fig. 4D). Specialists are faster and can better identify a transition into their associated stress than generalists, but the variety of environments experienced by cells make having a specialist for every environment implausible. We postulate that generalists avoid this constraint by providing an indirect channel that responds not to the extracellular signals sensed by specialists (*24*, *25*) but to intracellular signals (*26*, *27*), such as changes the ratio of ATP to ADP, cAMP, uncharged tRNAs, and the availability of amino acids (*16*). By detecting physiological perturbations, generalists are responsive to broader categories of stress (Fig. S15) – defined by their intracellular effects – and are agnostic to the environment’s precise nature. Generalists are therefore necessarily slower than specialists because they must wait for the environment to modify intracellular biochemistry. Indeed, we conjecture that the stochastic pulsing of the generalists in constant environments (*20*) is a response to spontaneous fluctuations in intracellular physiology.

As environments become more complex, multiple transcription factors are needed to generate an internal representation. Pooling the data to consider environments with different states (Fig. 4E), the mutual information plateaus as the numbers of transcription factors increase, with four sufficing to generate ~80% of the information. This increase comes both from the distinct dynamics of the transcription factors, such as differences in timing (Fig. S16), and from decreasing the effects of biochemical stochasticity by averaging the multiple readouts.

Together, our results show that it is only through the dynamic activation of multiple transcription factors that cells can encode sufficient information to generate a downstream response specific to an environment (*15*, *28*). Previous estimates focused on a single protein and found that information of one bit is typical (*5, 9, 29, 30*), implying that a cell can only distinguish between two or three environmental states. Collectively, transcription factors can complement each other through the diversity in their responses (*31*) as well as averaging biochemical stochasticity. Paralleling discoveries in neuroscience, we expect that such multi-dimensional internal representations are widespread within cellular biology and their failures in encoding information, by causing dysfunctional decision-making, can instigate deleterious behaviors and disease (*32*).

## Materials and Methods

### Time-lapse microscopy

BY strains with fluorescently-tagged transcription factors (*33*) were grown in ALCATRAS microfluidic devices (*21*) in synthetic complete medium with 2% glucose for at least 3 hours and then exposed to stress for another 5 hours (SI Appendix). Bright-field and fluorescence images were acquired every 2.5 min and cells were segmented using the DISCO algorithm (*34*).

### Estimating the mutual information

Our algorithm for estimating mutual information by decoding (SI Appendix) involves: (i) using principal component analysis to delineate a basis set for the time series; (ii) training a linear support vector machine to classify time series in this basis set; (iii) calculating a confusion matrix using the test data; (iv) estimating (a lower bound on) the mutual information by interpreting the entries of the confusion matrix as conditional probabilities.

## Statistics

All statistical analyses were performed in Matlab R2014b.

## Data

Available at http://dx.doi.org/10.7488/ds/2214

## Acknowledgments

We thank S. Jaramillo-Riveri, P. Thomas, M. Voliotis, E. Wallace, and the Swain laboratory for critical comments; I. Farquhar for preparing strains; and our funders: the BBSRC (JMJP & PSS), the EPSRC (AAG), and the Austrian Science Fund – FWF P28844 (GT).

## References

1. Tkačik G, Bialek W (2016) Information processing in living systems. Annu Rev Condens Matter Phys 7:89–117.

2. Gold JI, Shadlen MN (2007) The neural basis of decision making. Annu Rev Neurosci 30:535–574.

3. Perkins TJ, Swain PS (2009) Strategies for cellular decision-making. Mol Syst Biol 5:326.

4. Cai L, Dalal CK, Elowitz MB (2008) Frequency-modulated nuclear localization bursts coordinate gene regulation. Nature 455(7212):485–490.

5. Cheong R, Rhee A, Wang CJ, Nemenman I, Levchenko A (2011) Information transduction capacity of noisy biochemical signaling networks. Science 334(6054):354–358.

6. Hao N, O’Shea EK (2011) Signal-dependent dynamics of transcription factor translocation controls gene expression. Nat Struct Mol Biol 19(1):31–39.

7. Hao N, Budnik BA, Gunawardena J, O’Shea EK (2013) Tunable signal processing through modular control of transcription factor translocation. Science 339(6118):460–464.

8. Purvis JE, Lahav G (2013) Encoding and decoding cellular information through signaling dynamics. Cell 152(5):945–956.

9. Selimkhanov J, et al. (2014) Systems biology. Accurate information transmission through dynamic biochemical signaling networks. Science 346(6215):1370–1373.

10. Goulev Y, et al. (2017) Nonlinear feedback drives homeostatic plasticity in H2O2 stress response. eLife 6:e23971.

11. Elowitz MB, Levine AJ, Siggia ED, Swain PS (2002) Stochastic gene expression in a single cell. Science 297(5584):1183–1186.

12. López-Maury L, Marguerat S, Bähler J (2008) Tuning gene expression to changing environments: from rapid responses to evolutionary adaptation. Nat Rev Genet 9(8):583–593.

13. Mitchell A, Wei P, Lim WA (2015) Oscillatory stress stimulation uncovers an achilles heel of the yeast mapk signaling network. Science 350(6266):1379–1383.

14. Granados AA, et al. (2017) Distributing tasks via multiple input pathways increases cellular survival in stress. Elife 6:e21415.

15. Gasch AP, et al. (2000) Genomic expression programs in the response of yeast cells to environmental changes. Mol Biol Cell 11(12):4241–4257.

16. Broach JR (2012) Nutritional control of growth and development in yeast. Genetics 192(1):73–105.

17. Purvis JE, et al. (2012) p53 dynamics control cell fate. Science 336(6087):1440–1444.

18. Ashall L, et al. (2009) Pulsatile stimulation determines timing and specificity of NF-*κ*B-dependent transcription. Science 324(5924):242–246.

19. Shannon CE, Weaver W (1999) The mathematical theory of communication. (University of Illinois Press, Urbana, Illinois).

20. Dalal CK, Cai L, Lin Y, Rahbar K, Elowitz MB (2014) Pulsatile dynamics in the yeast proteome. Curr Biol 24(18):2189–2194.

21. Crane MM, Clark IBN, Bakker E, Smith S, Swain PS (2014) A microfluidic system for studying ageing and dynamic single-cell responses in budding yeast. PLoS ONE 9(6):e100042.

22. MacKay DJC (2003) Information theory, inference, and learning algorithms. (Oxford University Press, Oxford, U.K.).

23. Sharifpoor S, et al. (2011) A quantitative literature-curated gold standard for kinase-substrate pairs. Genome Biol 12(4):R39.

24. Delaunay A, Isnard AD, Toledano MB (2000) H2O2 sensing through oxidation of the Yap1 transcription factor. EMBO J 19(19):5157–5166.

25. Reiser V, Raitt DC, Saito H (2003) Yeast osmosensor Sln1 and plant cytokinin receptor Cre1 respond to changes in turgor pressure. J Cell Biol 161(6):1035–1040.

26. Dechant R, Saad S, Ibáñez AJ, Peter M (2014) Cytosolic pH regulates cell growth through distinct GTPases, Arf1 and Gtr1, to promote Ras/PKA and TORC1 activity. Mol Cell 55(3):409–421.

27. Filteau M, et al. (2015) Systematic identification of signal integration by protein kinase A. Proc Natl Acad Sci U S A 112(14):4501–4506.

28. Hansen AS, O’Shea EK (2013) Promoter decoding of transcription factor dynamics involves a trade-off between noise and control of gene expression. Mol Syst Biol 9:704.

29. Voliotis M, Perrett RM, McWilliams C, McArdle CA, Bowsher CG (2014) Information transfer by leaky, heterogeneous, protein kinase signaling systems. Proc Natl Acad Sci U S A 111(3):E326–E333.

30. Uda S, et al. (2013) Robustness and compensation of information transmission of signaling pathways. Science 341(6145):558–561.

31. Dubuis JO, Tkacik G, Wieschaus EF, Gregor T, Bialek W (2013) Positional information, in bits. Proc Natl Acad Sci U S A 110(41):16301–16308.

32. Luo Q, Beaver JM, Liu Y, Zhang Z (2017) Dynamics of p 53: A master decider of cell fate. Genes 8(2):66.

33. Huh WK, et al. (2003) Global analysis of protein localization in budding yeast. Nature 425(6959):686–691.

34. Bakker E, Swain PS, Crane MM (2017) Morphologically constrained and data informed cell segmentation of budding yeast. Bioinformatics in press.

